# Fast, flexible analysis of differences in cellular composition with crumblr

**DOI:** 10.1101/2025.01.29.635498

**Authors:** Gabriel E. Hoffman, Panos Roussos

**Author notes:** Correspondence to: G.E.H and P.R.

## Abstract

Changes in cell type composition play an important role in human health and disease. Recent advances in single-cell technology have enabled the measurement of cell type composition at increasing cell lineage resolution across large cohorts of individuals. Yet this raises new challenges for statistical analysis of these compositional data to identify changes in cell type frequency. We introduce `crumblr` (DiseaseNeurogenomics.github.io/crumblr), a scalable statistical method for analyzing count ratio data using precision-weighted linear mixed models incorporating random effects for complex study designs. Uniquely, `crumblr` performs statistical testing at multiple levels of the cell lineage hierarchy using a multivariate approach to increase power over tests of one cell type. In simulations, `crumblr` increases power compared to existing methods while controlling the false positive rate. We demonstrate the application of `crumblr` to published single-cell RNA-seq datasets for aging, tuberculosis infection in T cells, bone metastases from prostate cancer, and SARS-CoV-2 infection.

---

Tissues are composed of diverse cell types that play essential roles in human biology and disease ^1^. Advances in single-cell technologies have enabled the measurement of genome-wide gene expression profiles from millions of single cells across hundreds of samples ^2^. This has driven an improved understanding of gene expression dynamics within and across cell types, developmental stages, and disease states ^3,4^. Cellular composition is also dynamic, and single-cell technologies have enabled studies of the changes in the frequency of different cell types due to early developmental stage, adult aging, immune response, and disease progression ^5–9^.

As the scale of single-cell datasets continues to increase, study designs have become more complex, and the cell type resolution is expanding to consider lower-frequency cell types. Statistical tools should model these complex datasets, have the power to identify differences in cellular composition across samples, and control the false positive rate. Cell type frequency can vary due to biological factors of interest, but also due to tissue dissection, specimen quality and technical factors. In a typical workflow, data-derived cell clusters are identified based on gene expression profiles, and the counts for each cell cluster are then analyzed with a regression model to test for changes in frequency across samples based on a variable of interest (i.e., disease state) while accounting for confounding variables.

Analyses take one of two approaches to examine changes in cellular composition, and the choice of approach affects statistical power, handling of complex study designs, integration with downstream analyses, and computational scaling. Regression models either use cell fractions (i.e., frequency) or model cell counts directly. While using cell fractions is simplest, computing a fraction from cell counts loses valuable information about the precision of the measurement. Consider two samples with an equal fraction of neuronal cells, but the first observation counts 1000 neurons out of 5000 total cells, while the second counts one neuron out of 5 total cells. Of course, 1000 / 5000 is a more precise measurement than 1 / 5 while having the same cell fraction. Similarly, measurements of lower-frequency cell types are less precise when the total number of cells observed is the same. Translating this intuition and lessons learned from modeling counts in RNA-seq data ^10,11^ to the case of compositional data analysis, modeling the variation in measurement precision is essential to maximize statistical power when the total number of counts or the cell frequency varies widely across samples. Both of these cases are common in single-cell datasets.

The most widely used methods that analyze cell fractions are simple linear models using either observed cell fractions or some transformation of these fractions as the regression response. Linear models assume normally distributed errors, and this assumption is better satisfied after transforming the cell fractions using a log, logit, arcsin-sqrt or the centered log ratio (CLR) transforms ^12,13^. These methods are fast but sacrifice statistical power because they don’t consider the precision of the cell fraction measurements. In addition, cell fractions can also be modeled directly with binomial or beta-binomial models to address this issue partially, but these also ignore the magnitude of the counts. Alternatively, methods that directly use count data and thus model variation in measurement precision include Poisson and negative binomial models, as well as the hierarchical Bayesian model of scCODA ^14^. Yet these methods may not control the false positive rate across all conditions, can be computationally demanding, and don’t easily integrate into other downstream analyses.

Here, we introduce count ratio uncertainty modeling based linear regression (`crumblr`) for differential analysis of cellular composition that models variation in measurement precision, handles complex study designs with random effects, and performs tests at multiple levels of the cell lineage hierarchy using a multivariate approach. The `crumblr` framework has high statistical power, controls the false positive rate while being scalable to large single-cell datasets, and integrates with widely used R/Bioconductor packages.

## Results

### crumblr workflow for differential cellular composition analysis

The `crumblr` framework enables analysis of compositional data starting from cell cluster counts by transforming the count data and using weighted regression models for variance partitioning analysis, differential composition testing with univariate tests, and multivariate testing along a cellular hierarchy (**Fig 1A**). The `crumblr` approach models observed cell counts following transformation with the centered log ratio (CLR). The CLR transform is widely used in compositional data analysis and normalizes each cell component with the same denominator using the geometric mean of cell frequencies ^12,15,16^. The CLR can be evaluated in log space as a linear combination (i.e., weighted sum) of the log proportions and transforms fractions for use as responses or covariates in regression models, as well as PCA and hierarchical clustering ^12,15,16^. CLR can naturally be used in linear mixed models, allowing for random effects and multivariate testing.

**Figure 1:**
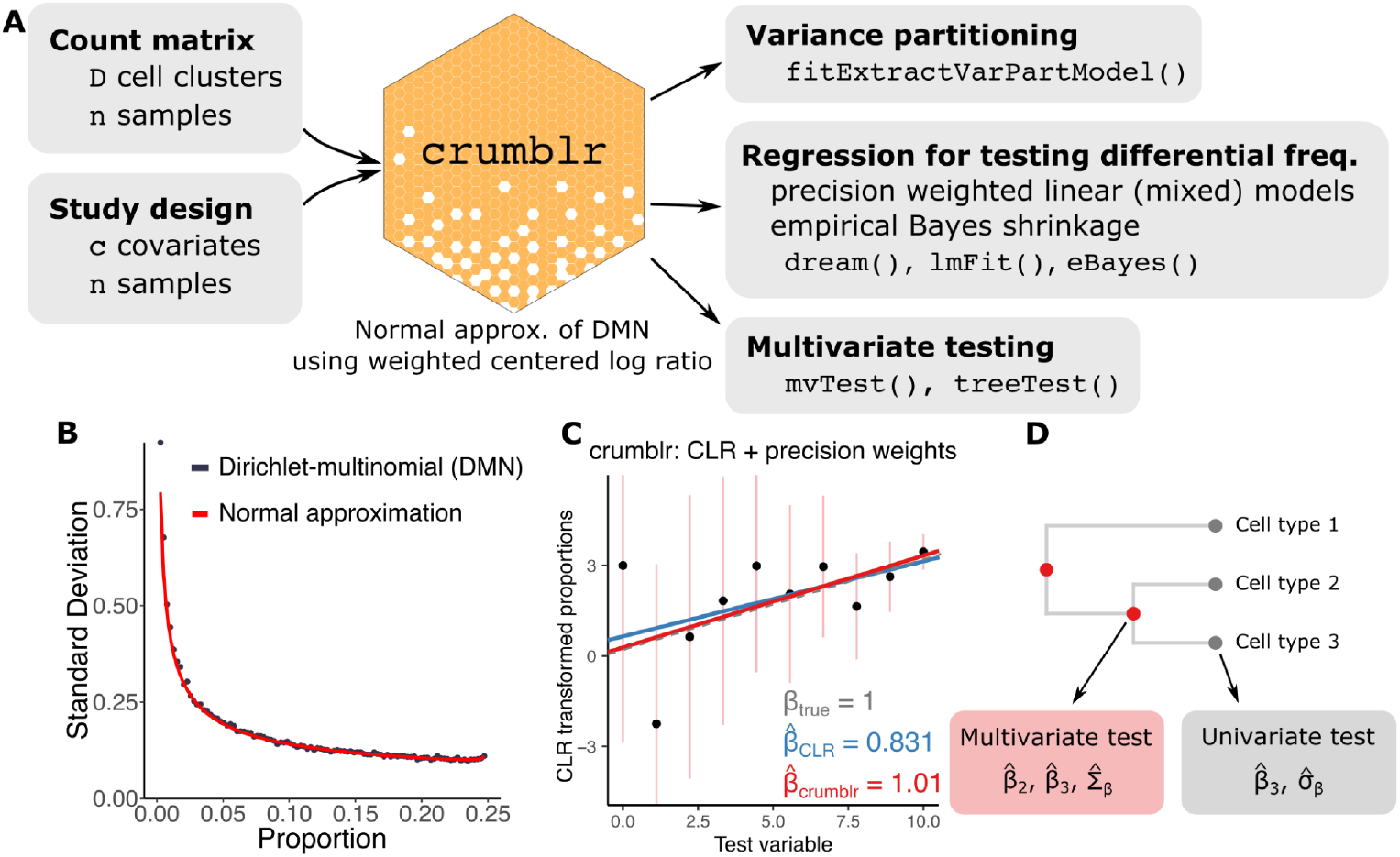
crumblr analysis workflow. **A)** `crumblr` transformation of count matrix enables precision weighting for variance partition analysis, linear mixed model testing for differential cell frequency, and multivariate testing along a cellular hierarchy. **B)** Standard deviation of the CLR transformed proportions from 1000 samples drawn from a Dirichlet-multinomial distribution (blue points) are compared to the normal approximation developed here (red line) when at least two counts are observed. Data was simulated using an overdispersion of τ = 10, 500 total counts across 15 categories and 1000 simulations. **C)** Illustration showing standard regression fit on CLR transformed proportions (blue line), and precision-weighted regression using `crumblr` (red line) that models the measurement uncertainty in the count ratios, shown here by confidence intervals. **D)** Multivariate hypothesis testing in `crumblr` enables analysis of internal nodes in a hierarchical clustering of cell types while modeling estimated effect sizes and the correlation between them.

Yet CLR transforms the cell *fractions* and does not consider the observed *counts* or the variation in measurement precision. The `crumblr` framework applies the CLR transform and uses an asymptotic normal approximation of the Dirichlet-multinomial distribution to estimate the sampling variance of the transformed fractions (see **Methods**). This approximation works well across a wide range of cell counts and proportions (**Fig 1B**). These sample variances are used in precision-weighted regressions to model variation in measurement precision (**Fig 1C**), as is widely used in differential expression analysis ^10,17^. Transforming compositional data analysis into a precision-weighted regression model enables the incorporation of random effects and empirical Bayes shrinkage of test statistics. In addition to univariate testing of each cell cluster, precision-weighted models enable testing at multiple levels of the cell lineage hierarchy using a multivariate approach to increase power over tests of one cell cluster. Compared to 10 other methods for testing changes in cellular composition, only `crumblr` has high statistical power, controls the false positive rate across conditions, and offers flexibility to incorporate random effects and multivariate testing while being scalable to large datasets (**Fig S1**). The `crumblr` workflow easily integrates with widely used R/Bioconductor packages.

### Performance on simulated data

We evaluated the performance of `crumblr` and 10 other methods to identify differences in cell frequencies in simulations across a broad range of conditions. Count data was drawn from a Dirichlet-multinomial distribution, and baseline simulations were performed with 200 samples, a variable of interest drawn from a standard normal distribution, 10 cell clusters, of which one had differential frequency with an effect size of 0.2, a mean of 2000 cells observed for each sample, no batch effect, overdispersion parameter of 10, and 10K simulated datasets. The `crumblr` and negative binomial models showed the highest area under the precision-recall (AUPR) curve to identify the cell clusters with a change in frequency driven by the simulated variable of interest (**Fig 2A**). Methods that did not model the measurement precision (i.e., linear regression on fraction, log fraction, logit fraction, arcsin-sqrt fraction, or CLR) had lower power, and methods that did not model overdispersion of count data (i.e., Poisson and binomial models) did not control the false positive rate. Further simulations modified the parameters of this baseline simulation by varying the total cell counts between 100 and 4000 (**Fig S2**), number of cell clusters between 7 and 20 (**Fig S3**), sample size between 20 and 200 (**Fig S4**), count overdispersion parameter between 1 and 20 (**Fig S5**), and variance explained by batch effect between 0 and 20% (**Fig S6**). Across all these conditions, `crumblr` and the negative binomial model consistently showed the highest power, while `crumblr` controlled the false positive rate even for small sample sizes. Both `crumblr` and the negative binomial model were fast and required < 10 seconds to analyze 500 samples with 20 cell clusters (**Fig S7**). Rigorous analysis with scCODA ^14^ was limited by its long run time (∼1000 seconds on this dataset), and evaluation on 100 simulated datasets showed lower statistical power than `crumblr` using default settings (**Fig S8**).

**Figure 2:**
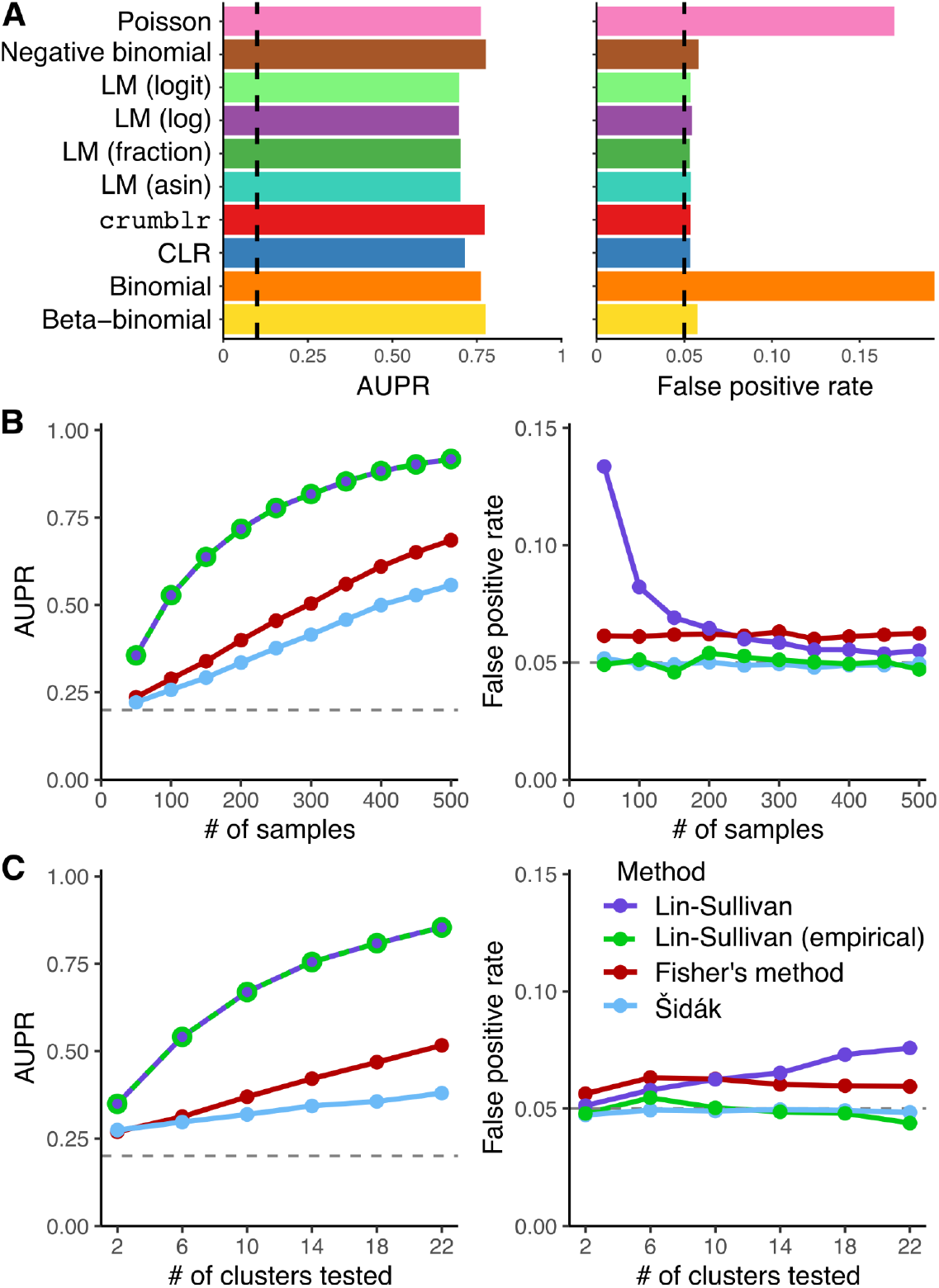
Performance on simulated cellar composition data. **A)** Performance of univariate tests across 10k simulated datasets under baseline parameters. For linear models indicated with ‘LM’, ‘fraction’ indicates using the cell fraction, *f*, as the response, ‘log’ indicates *log* (*f*), ‘logit’ indicates *logit*(*f*) = *log*(*f*) - *log*(*l* – *f*), and ‘asin’ indicates 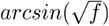. The area under the precision-recall curve (AUPR) (left) and false positive rate (FPR) at a 5% cutoff computed under the null model (right). For AUPR, the dashed line indicates the performance of the random method. For FPR, the dashed line indicates the target for a properly calibrated method. **B)** Performance of multivariate testing in simulated data for increasing sample sizes. AUPR (left) and FPR (right) of 4 methods for multivariate testing. **C)** Performance of multivariate testing in simulated data for increasing number of cell clusters tested. AUPR (left) and FPR (right).

In addition to performing univariate testing of each cell cluster individually, `crumblr` can perform multivariate testing along the cell lineage hierarchy to test for changes in cellular composition at multiple resolutions. In a hierarchy, testing an internal node with *m* children corresponds to a multivariate test of the *m* child cell clusters. We evaluated the performance of 3 existing methods to perform multivariate testing by combining the results for univariate tests from `crumblr` . Based on these simulations varying the number of samples (**Fig 2B**) and the cell clusters *m* (**Fig 2C**), we developed a novel statistical test with high power that controls the false positive rate.

Of these multivariate methods, the Šidák method is the simplest and reports the smallest p-value corrected for the number of clusters tested. This method had low power because it uses only the smallest p-value. Fisher’s method combines p-values from all tests and compares a test statistic to a chi-squared null distribution. Yet these methods don’t consider the effect size and assume that all tests are independent, giving elevated false positive rates. Instead, we use a fixed effect meta-analysis of the estimated effect size while modeling the correlation in the estimates using the Lin-Sullivan test ^18^. This approach substantially increases the power to find sets of cell clusters with differential frequency, but for small sample sizes it gave an elevated false positive rate since it depends on an asymptotic null distribution. We developed an extension to the Lin-Sullivan test using an empirical null distribution that controls the false positive rate for small sample sizes while retaining high power (see **Methods**).

### Cell type composition changes with age in PBMCs

The biology of the human immune system depends on age, with changes in both gene expression programs and cellular composition over time ^19^. However, a detailed understanding of the dynamics of high-resolution cell types has been limited by the ability to measure cellular composition across a cohort with a broad age range. Yazar, et al. ^20^ generated single-cell transcriptome data for 1.2M peripheral blood mononuclear cells (PBMCs) from 982 donors in the OneK1K cohort with ages ranging from 19 to 97 years, and identified in 29 data-derived clusters with 23 having at least 1000 cells. Variance partitioning analysis showed substantial variation in cell type frequency across the 75 batches, but little variation between males and females (**Fig. 3A**). Age explained 35.9% of the variation in frequency for naive CD8^+^ αβ T cells and greater than 5% for five other cell clusters. Univariate analysis identified the strongest decreases in frequency of CD8^+^ αβ T cells (β = -0.0297, p = 3.6e-93, FDR < 1e-10) and mucosal invariant T cells (β = -0.0120, p = 2.8e-14, FDR = < 1e-10) with age, along with significant decreases in 2 additional clusters and significant increases in 10 clusters (**Fig. 3B**). Hierarchical clustering of gene expression profiles and multivariate testing of cellular composition shows widespread changes in immune cell frequency throughout human aging (**Fig. 3C**). Examining the trajectory of CD8^+^ αβ T cell frequency shows a marked decrease over age, and also highlights the importance of modeling the measurement error with `crumblr` (**Fig. 3D**). Since the frequency of CD8^+^ αβ T cells decreases with age, measurement precision also decreases with age. While the `crumblr` model supports a linear trend between transformed frequency and age, a model ignoring the varying measurement precision supports a non-linear trend with an accelerated decrease in frequency after age ∼70 (p = 5.20e-3).

**Figure 3:**
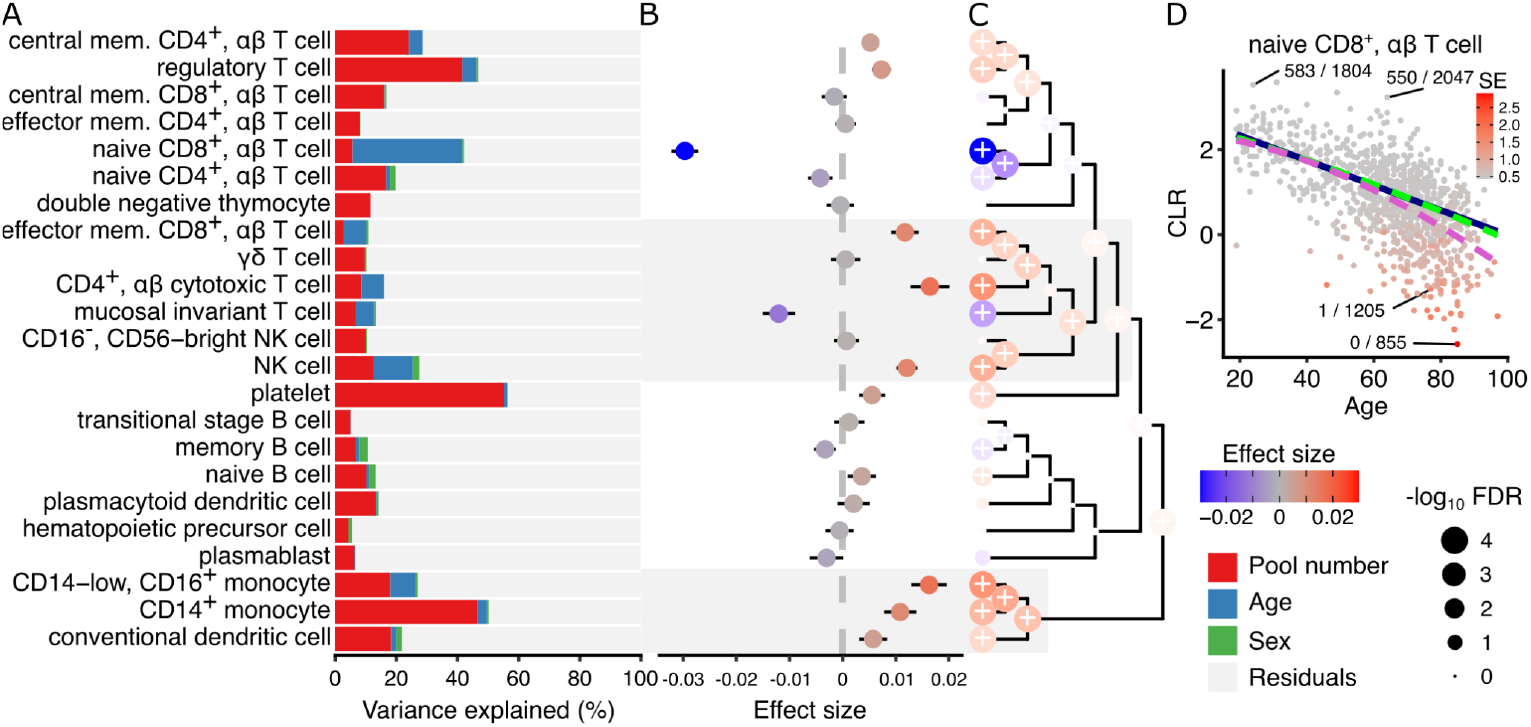
crumblr identifies compositional changes associated with aging in blood. **A)** Variance partitioning analysis quantifies the contribution of each variable to variation in frequency for each cell type. **B)** Estimated effect size from regression analysis of composition associated with donor age. A positive coefficient estimate indicates increased cell type frequency with age. The color indicates effect size, and the bars indicate 95% confidence interval. **C)** Hierarchical clustering of cell types with colored points indicating estimated effect sizes for each cell type at leaves and multivariate testing at internal nodes. Results on leaves match results in (B). Point size indicates FDR, and ‘+’ indicates FDR < 5%. **D)** Scatter plot of CLR-transformed naive CD8^+^ αβ T cell frequency versus donor age. A point indicates a single donor and the color indicates the standard error from limited measurement precision. A subset of points are labeled with the observed cell type fraction with, for example, 1 / 1205 indicating 1 CD8^+^ αβ T cell out of 1205 total cells. Regression trends are shown for a linear fit (blue) and quadratic fit (green) modeling the varying measurement precision using `crumblr`, and a quadratic fit (purple) ignoring measurement precision.

### Decrease in CD4^+^ T_h_1 and T_h_17 T cells following tuberculosis infection

Immune response to infection can change the cellular composition at multiple levels. Nathan, et al. ^21^ studied the post-disease steady state of T cells following tuberculosis (TB) infection and disease resolution, by isolating 500K T cells from 259 donors, including 128 who had previously had TB. After isolating T cells from PBMCs and applying CITE-seq to measure gene expression and surface proteins for each cell, their analysis identified 31 data-derived T cell clusters. The average frequency of these T cell clusters ranged from 0.062 to 8.6%, with the frequency of each cluster varying by over 10-fold across individuals (**Fig S9**). The original analysis found that the frequency of some clusters correlated with age, sex, ancestry, and the season of the blood draw and identified a cluster of CD4^+^ T_h_17 T cells showing a significant decrease in frequency in individuals with prior TB infection.

We applied `crumblr` to identify changes in cell type composition associated with TB status for each of the 31 data-derived T cell clusters, as well as hierarchical clusters. Variance partitioning analysis showed the substantial contribution of age to T cell composition, while season and sex explained less variation (**Fig. 4A**). Univariate analysis of differential cell frequency based on TB status identified the same CD4^+^ T_h_17 cluster as having the strongest estimated effect (β = -0.226, p = 3.73e-6, FDR = 0.00011), but also identified three other T_h_1 or T_h_17 clusters with significant decreases in TB-infected individuals (**Fig. 4B**). In addition, CD4^+^ cytotoxic T cells show the largest increases in frequency. Hierarchical clustering of the annotated cell types based on gene expression placed these the 4 T_h_1 / T_h_17 cell clusters together, and multivariate differential frequency analysis showed that the parent node of these cell types had significantly decreased frequency following TB infection (β = -0.152, p = 3.04e-6, FDR = 1.06e-4) (**Fig. 4C**). Highlighting the CD4^+^ T_h_17 cluster shows the decrease in frequency following TB infection, and also shows that samples with low frequency of this cell type also have high measurement error (**Fig. 4D**).

**Figure 4:**
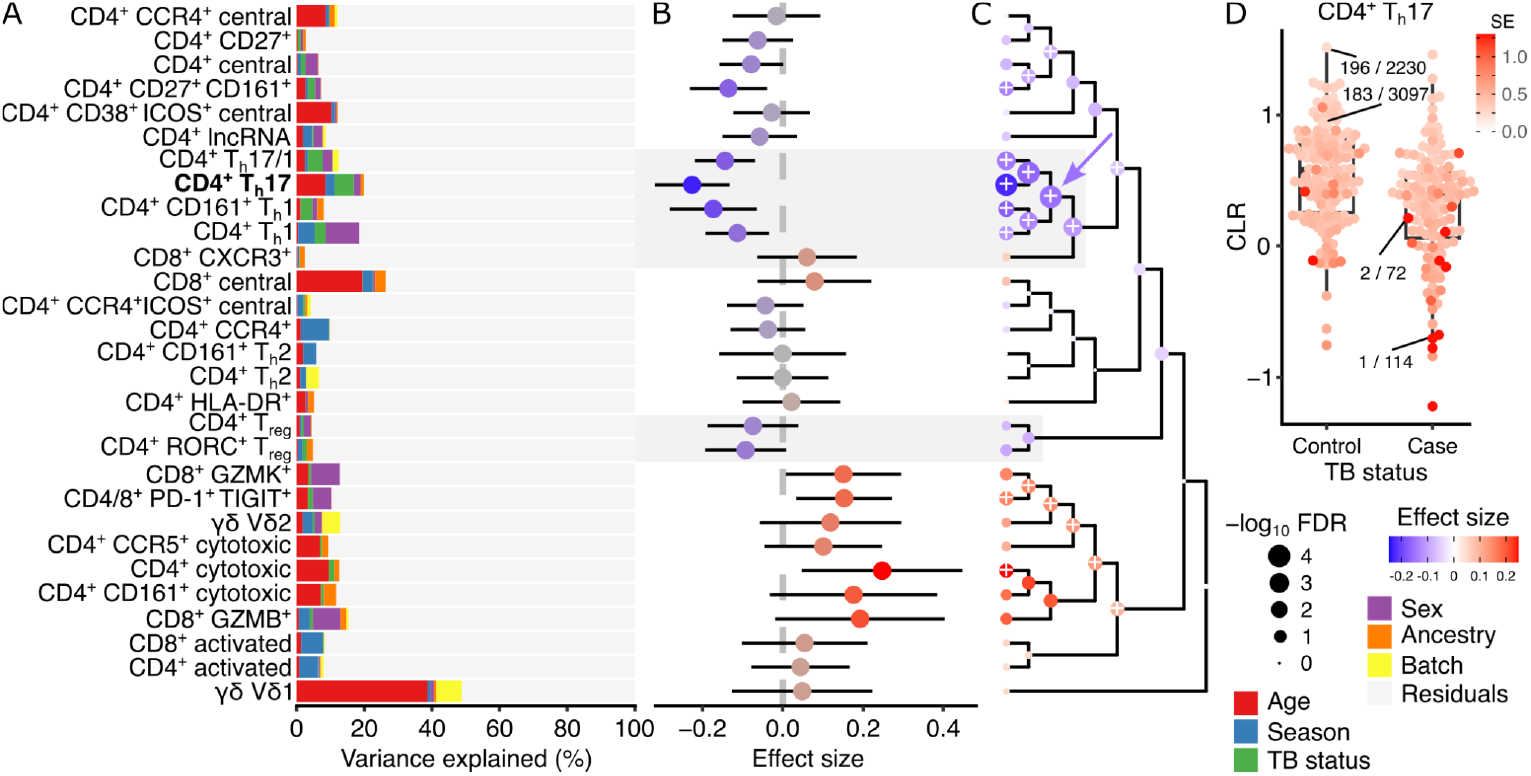
crumblr identifies compositional changes associated with tuberculosis infection in T cell subpopulations in 259 donors. **A)** Variance partitioning analysis quantifies the contribution of each variable to variation in frequency for each cell cluster. **B)** Estimated effect size by comparing the composition of TB-inflected and controls. A positive coefficient estimate indicates an increase in the frequency of TB infection. Color indicates effect size, and bars indicate a 95% confidence interval. **C)** Hierarchical clustering of cell types with colored points indicating estimated effect sizes for each cell type at leaves and multivariate testing at internal nodes. Results on leaves match results in (B). Point size indicates FDR, and ‘+’ indicates FDR < 5%. The parent node mentioned in the text is indicated by the blue arrow. **D)** Transformed frequency of CD4^+^ T_h_17 T cells in healthy controls and donors with previous TB infection. A point indicates a single donor, and the color indicates the standard error from limited measurement precision using `crumblr` . A subset of points are labeled with the observed cell type fraction with, for example, 1 / 144, indicating 1 CD4^+^ T_h_17 T cell out of 114 total cells.

### Cell type composition in bone metastases from prostate cancer

Bone metastases from prostate cancer result in poor patient prognosis ^22^. Single-cell transcriptomics has been applied to solid tumors, involved bone marrow, and distal bone marrow from 9 prostate cancer patients with bone metastases, as well as benign bone marrow from 7 patients without cancer to study changes in gene expression and cell composition ^23^. We applied `crumblr` to identify compositional differences associated with disease status. Variance partitioning analysis shows substantial variation in cell composition across patients, disease status of each sample (i.e., tumor, involved bone marrow, etc.), and disease status of the patient (i.e., cancer vs non-cancer) (**Fig. 5A**). Univariate analysis of composition differences between solid tumor and involved bone marrow identified significant changes in 10 of the 28 cell types (**Fig. 5B**). A subset of monocytes showed the strongest decrease in frequency, while pericytes, osteoblasts and endothelial cells showed the strongest increase. Hierarchical clustering of gene expression profiles and multivariate testing identified the parent node of the three monocyte subsets as having significantly decreased frequency (β = -1.10, p = 9.28e-4, FDR = 3.93e-3), and the parent node of pericytes, osteoblasts and endothelial cells having significantly increased frequency (β = 2.74, p = 9.15e-0, FDR = 3.92e-3) (**Fig. 5C**). Examining pericytes across the four disease states shows a substantial increase in tumor samples (**Fig. 5D**).

**Figure 5:**
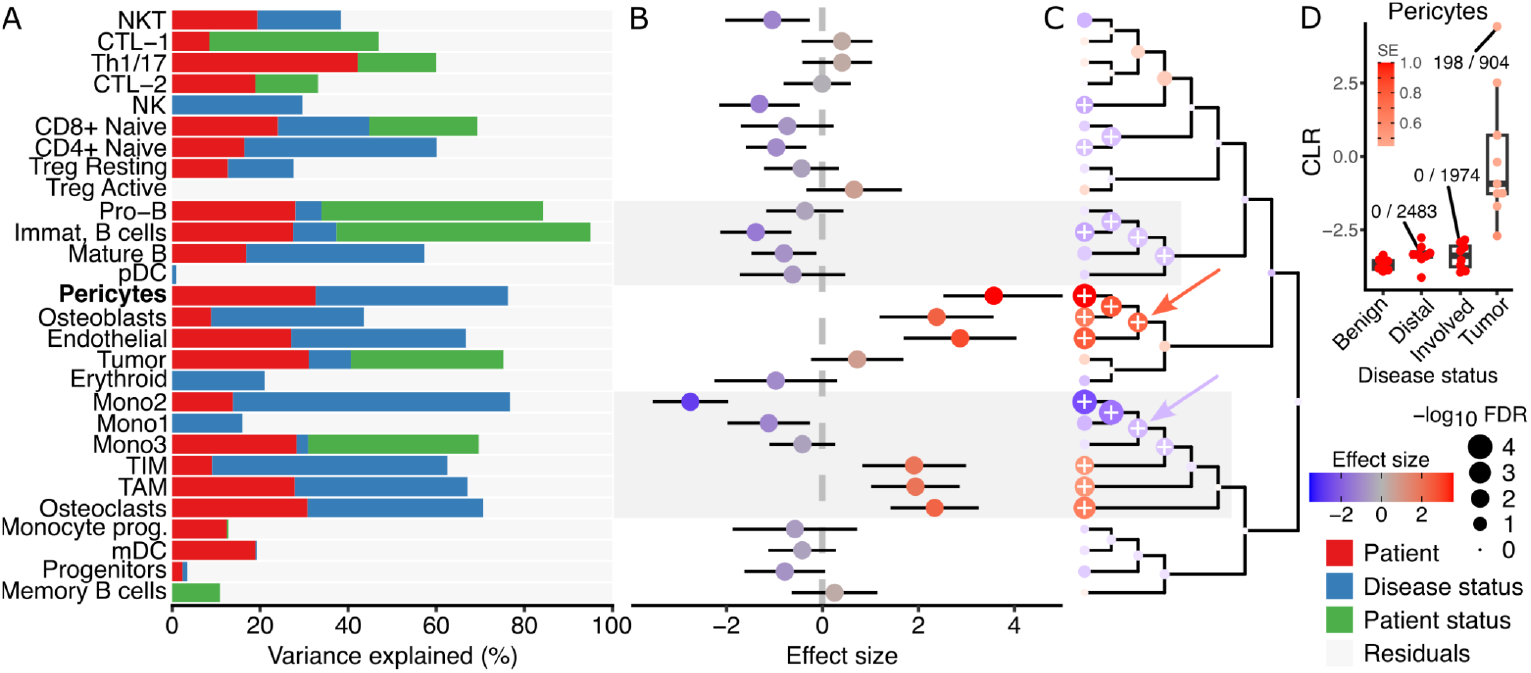
crumblr identifies composition changes in bone metastases from prostate cancer. **A)** Variance partitioning analysis quantifies the contribution of each variable to variation in frequency for each cell type. **B)** Estimated effect size by comparing the composition of solid tumors and involved bone marrow. The color indicates effect size, and the bars indicate a 95% confidence interval. **C)** Hierarchical clustering of cell types with colored points indicating estimated effect sizes for each cell type at leaves and multivariate testing at internal nodes. Results on leaves match results in (B). Point size indicates FDR, and ‘+’ indicates FDR < 5%. Parent nodes mentioned in the text are indicated by red and blue arrows. **D)** Transformed frequency of pericytes for each disease state. Each point indicates a single sample, and the color indicates the standard error from limited measurement precision using `crumblr` . A subset of points are labeled with the observed cell type fraction with, for example, 198 / 904, indicating 198 pericytes out of 904 total cells.

### Compositional changes associated with infection response

Viral and bacterial infections trigger an immune response, causing changes in gene expression and cell composition in blood. In order to study immune response to SARS-CoV-2 infection, the COMBAT Consortium ^24^ collected blood from patients hospitalized with COVID-19, along with blood from non-hospitalized COVID-19 patients, healthy controls, and patients with sepsis or flu. They compiled a single-cell transcriptome resource of 787K cells from 121 donors and identified 40 data-derived cell clusters. We applied `crumblr` to identify compositional differences associated with disease status. Variance partitioning analysis identified disease status as a major source of compositional variation, explaining more than 10% variance in 18 cell clusters (**Fig 6A**). Notably, age explains 26.4% of the variance in CD8^+^ naive T cells, and this cluster decreases in frequency with age (β =-0.271, p = 6.2e-10, FDR = 2.5e-8). Univariate analysis of the compositional difference between patients with severe COVID and healthy controls identifies significant decreases in 6 cell clusters, with the largest decreases in MAIT, plasmacytoid dendritic cell, invariant natural killer T cells (**Fig 6B**). Meanwhile, eight cell clusters increased in frequency, with the largest seen in plasmablasts, platelets, and cycling classical monocytes. Hierarchical clustering using gene expression profiles and multivariate testing shows limited higher-order changes in cell type frequency (**Fig 6C**). Examining classical cycling monocytes shows a baseline frequency shared between health donors and COVID-19 patients who have not been hospitalized, while frequency increases with COVID-19 severity in hospitalized patients to a level equal to that in sepsis patients (**Fig 6D**). This increase in frequency with COVID-19 severity is observed across multiple cell types (**Fig 6E**).

**Figure 6:**
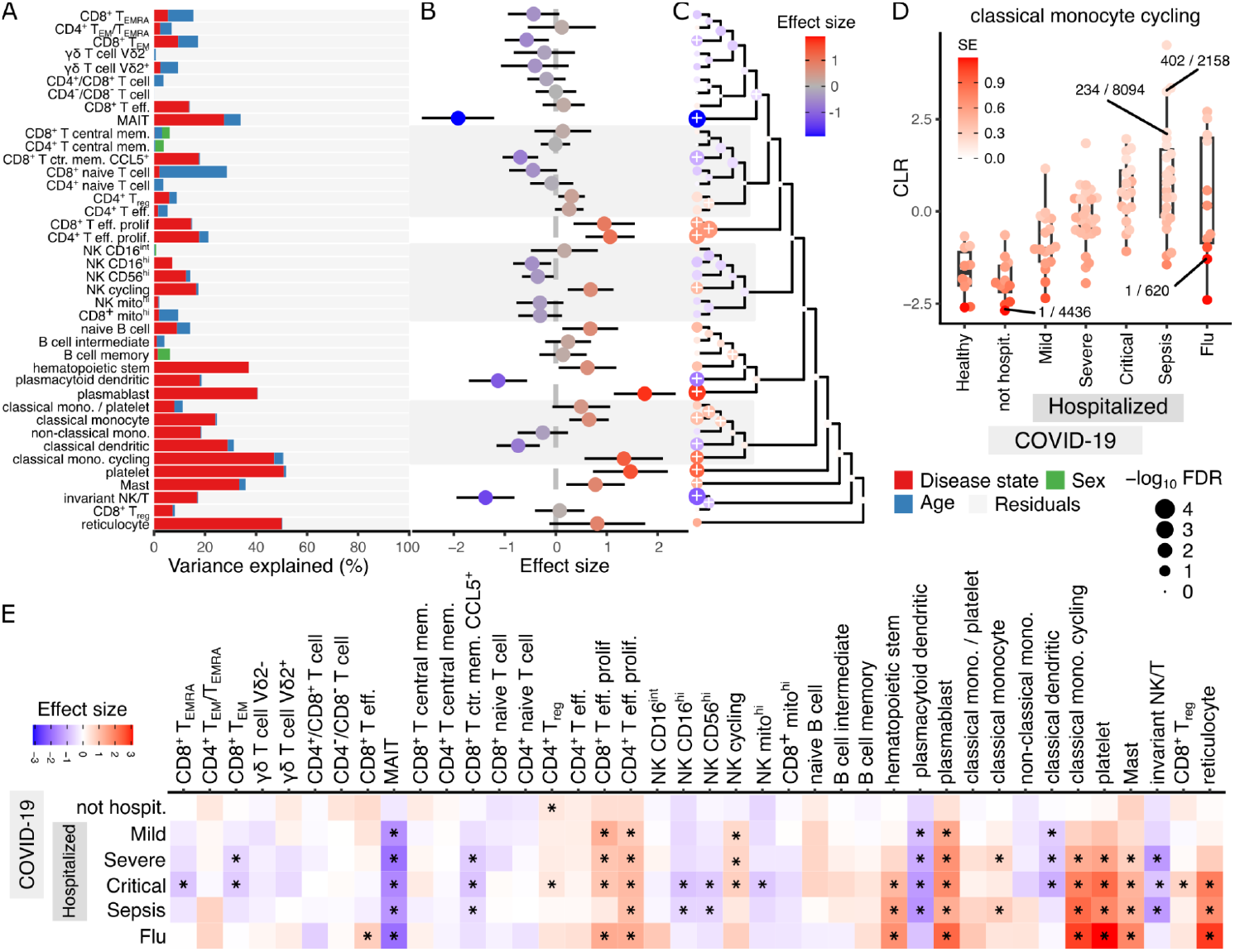
crumblr identifies compositional changes associated with infection. **A)** Variance partitioning analysis quantifies the contribution of each variable to variation in frequency for each cell type. **B)** Estimated effect size corresponding to leaves in (A) comparing composition between patients with severe COVID and healthy controls. A positive coefficient estimate indicates an increase in cell type frequency with infection. The color indicates effect size, and the bars indicate a 95% confidence interval. **C)** Hierarchical clustering of cell types with colored points indicating estimated effect sizes for each cell type at leaves and multivariate testing at internal nodes. Results on leaves match results in (B). Point size indicates FDR, and ‘+’ indicates FDR < 5%. **D)** Transformed frequency of cycling classical monocytes for each disease state. Each point indicates a single donor and the color indicates the standard error from limited measurement precision using `crumblr` . A subset of points are labeled with the observed cell type fraction with, for example, 1 / 620 indicating 1 cycling classical monocyte out of 620 total cells. **E)** Effect size estimates comparing cell type frequency to healthy controls. Color indicates effect size, and ‘*’ indicates FDR < 5%.

## Discussion

Here, we present the `crumblr` statistical framework for differential analysis of cellular composition to identify cell types that change frequency with a variable of interest. Across a broad range of simulation conditions, `crumblr` achieves high power while controlling the false positive rate. In addition to univariate tests of differential frequency, `crumblr` performs statistical testing at multiple levels of the cell lineage hierarchy using multivariate regression to increase power over tests of one cell component. The `crumblr` framework is fast, scalable to large single cell datasets and integrates with existing R/Bioconductor workflows from dreamlet ^25^, variancePartition ^17,26^ and limma ^27^.

Applying `crumblr` to 4 published single cell datasets identified biologically important changes in cellular composition. Analysis of T cells after tuberculosis infection identified multiple clusters of CD4^+^ T_h_1 and T_h_17 cells that decrease in frequency. Analysis of bone metastases from prostate cancer identified a decrease in monocyte frequency, and analysis of SARS-CoV2 infection response identified an increase in cycling classical monocytes that tracked with COVID-19 severity. Finally, analysis of compositional changes in blood with aging in the OneK1K cohort identified a decrease in naive CD8^+^ αβ T cells and mucosal invariant T cells with age. This trend between frequency and age is also observed in the COMBAT cohort, and is consistent with recent work on dynamics of T cell frequency with age ^7,28^. Changes in the thymus over the human lifespan are known to result in decreased production of naive T cells with age ^29^.

Here, we demonstrated the importance of modeling variation in measurement precision in count ratios when analyzing changes in cell frequency. Based on statistical theory, simulations and observations from real data, we see that measurement precision increases with cell frequency and with the total number of cells observed. Modeling this variable precision is especially important when the total number of counts of the cell frequency varies widely across samples.

The open source `crumblr` package and documentation are available at DiseaseNeurogenomics.github.io/crumblr and Bioconductor (pending) and will enable powerful analysis of differences in cellular composition.

## Methods

### Centered log-ratio transform for compositional data

Let the vector P store the proportions for each of *D* cell types. Then, the CLR transform of the proportion for cell type *i* is

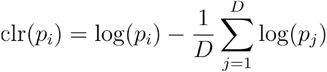

The CLR transform is widely used for compositional data analysis because it satisfies scale invariance, subcompositional coherence, invariance to selection of the category used as a reference, and produces unbounded values that can be well approximated by a normal distribution ^12,15^. Importantly, the CLR transform is a linear combination (i.e., weighted sum) of the log proportions. This property is central to the convenient form of the results described below.

Consider the observed proportions as random variables sampled from a Dirichlet-multinomial distribution with overdispersion parameter *τ*. As the number of counts, *n*, increases, the distribution of the estimated proportion converges to

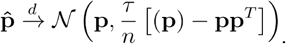

Setting *τ* = 1 corresponds to the multinomial distribution, and *τ* > 1 indicating overdispersion corresponds to the more general Dirichlet-multinomial distribution.

We apply the delta method to obtain an asymptotic normal approximation of the CLR transformed proportion under the Dirichlet-multinomial model (**Supplementary Methods**):

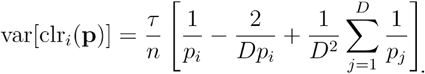

The properties of the CLR transform make this formula simpler than for other transformations ^12,30^.

Tests of the association are performed by fitting a linear or linear mixed model for each cell type using CLR transformed proportions as the response and the inverse variances as precision weights. Importantly, the estimated coefficient values and their covariance are invariant to the scaling of the precision weights. Since the overdispersion is multiplicative and the results are scaling invariant, the value *τ* of can be set to 1 instead of being estimated from the data. This property enables our framework to be widely applicable to cases of overdispersion, including cases where the overdispersion factor varies across components.

For real data, the asymptotic variance formula can give weights that vary substantially across samples and give very high weights for a subset of samples. In order to address this, we regularize the weights to reduce the variation in the weights to have a maximum ratio (default of 5) between the maximum and specified quantile value (default of 5%).

Regression models are fit separately across each of the *D* cell types. An empirical Bayes moderated t-statistic ^31^ is used by applying an inverse gamma prior to the residual variance for each cell type component. For linear mixed models, a recent extension is used to estimate the residual degrees of freedom for the empirical Bayes step ^17,25^.

### Multivariate testing combining multiple cell-type components

Consider a multivariate regression with *n* samples, *c* variables, and *m* response values using design matrix *X*_*n*×*c*_ and responses stored as columns in the matrix *Y*_*n*×*m*_. Standard results from multivariate regression with no weights (i.e., all samples have equal weights) give coefficient estimates

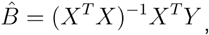

and variance-covariance matrix

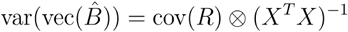

where 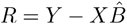 is the matrix of residuals, the vec(·) operator converts a matrix to a column vector, and ⊗ is the Kronecker product.

These coefficients and variance-covariance matrix can then be used in hypothesis testing across responses in the multivariate model. Here, we use a fixed effects meta-analysis to account for covariance between the coefficient estimates ^18^. Letting 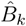 be the vector of coefficient estimates for variable *k* across *m* responses, 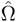 be the estimated covariance matrix, and be a vector of 1’s, the test statistic

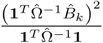

is asymptotically distributed as a 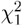 under the null. When 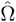 is diagonal, so there is no covariance between coefficient estimates; this reduces to standard fixed effects meta-analysis ^32^.

### Finite sample null distribution

The theoretical null distribution for the Lin-Sullivan statistic for fixed effects meta-analysis is asymptotically 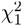, so computing the p-value of an observed test statistic using the 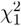 distribution works well when the regression coefficients are estimated from data with sufficient sample size. However, for a finite sample size, this null distribution is no longer 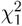, and this theoretical null can give an inflated false positive rate with a small sample size.

Instead, we sample test statistics from the null distribution based on a hierarchical model. We draw covariance matrices from a Wishart distribution, sample coefficients from a multivariate normal with this covariance, and then compute the Lin-Sullivan statistic. These Monte Carlo draws are sampled according to

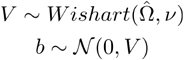

where *ν* is the degrees of freedom from the regression analysis. A gamma distribution is then fit to these draws from the null distribution, and an empirical p-value is computed from the cumulative distribution function of this gamma.

### Extension to weighted multivariate regression

When weights are not equal to 1 and vary for each response, the coefficient estimates and their covariance have a more complicated form. Letting *w*^(*i*)^ be the vector of weights for response *i, W*^(*i*)^ = diag (*w*^(*i*)^), and *y*_*i*_ be the vector storing response *i*, the coefficient estimates for response *i* are

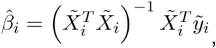

where 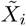 and 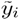 represent the weighted design and response according to 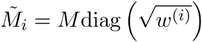 for any matrix or vector *M*. The covariance between coefficient estimates for responses *i* and *j* is

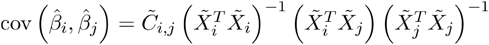

where 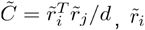, is the weight residuals for response, *i* and *d* is the residual degrees of freedom. When all weights are 1, this reduces to the standard results above.

### Extension to weighted linear mixed models

For the linear mixed model, closed forms don’t exist. Instead, the coefficients and variance-covariance matrix are estimated using numerical optimization ^33^. Letting 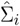 be the estimated variance-covariance matrix between coefficients for response *i*, the covariance between coefficients for responses and can be approximated as

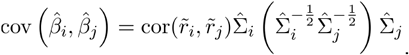

When this formula is applied to the fixed effect model, it is equivalent to the previous formula when the weights vary across samples but are shared across responses so that *w*^(*i*)^ and *w*^(*j*)^ are equal (**Supplementary Methods**).

### Comparison with other methods

Hypothesis testing for differential cell composition was performed with a range of existing methods that either modeled the cell fraction or directly modeled the number of observed counts (**Supplementary Methods**).

### Simulations of multivariate testing

In the first simulation, m=15 responses were used and sample size ranges from 50 to 500 with increments of 50. In the second simulation, 200 samples were simulated with m in 2 to 22 responses in increments of 4. In both cases, 100k simulations were performed using a design matrix including an intercept term and a variable sampled for a standard normal, an effect size of 0.25 shared across response, error covariance sampled for a multivariate normal using `clusterGeneration::genPositiveDefMat(m, ratioLambda= 100)` .

## Code Availability

The `crumblr` R package and documentation are available at DiseaseNeuroGenomics.github.io/crumblr and will soon be available at bioconductor.org/packages/crumblr (pending). The multivariate regression analysis proposed here is implemented in our `variancePartition` package using the function `mvTest()` to post-process results from `dream()` . Multivariate hypothesis testing is implemented in our `remaCor` package, which includes the novel extension of the Lin-Sullivan test for small sample sizes. Code for analysis and simulations is available at https://github.com/GabrielHoffman/crumblr_analysis.

### Software versions

`R v.4.4.1`

`crumblr v0.99.11`

`variancePartition v1.36.3`

`lme4 1.1.35.5`

`scCODA v0.1.9`

## Author Contributions

G.E.H. developed the `crumblr` package and performed analysis. G.E.H. and P.R. supervised the analysis and wrote the manuscript.

