## Supplementary Methods, Figures, Tables for "Fast, flexible analysis of differences in cellular composition with crumblr"

### Supplementary Figures

| Method | High power for univariate test | Controls false positive rate | Models varying measurement precision | Models random effects | Robust to overdispersion | Variance partitioning analysis | Multivariate testing | Fast (runtime $<$ 10s) |
|---:|---|---|---|---|---|---|---|---|
| `crumblr` | ✓ | ✓ | ✓ | ✓ | ✓ | ✓ | ✓ | ✓ |
| CLR | | ✓ | | ✓ | | ✓ | ✓ | ✓ |
| Linear Model (frac) | | ✓ | | ✓ | | ✓ | ✓ | ✓ |
| Linear Model (log) | | ✓ | | ✓ | | ✓ | ✓ | ✓ |
| Linear Model (logit) | | ✓ | | ✓ | | ✓ | ✓ | ✓ |
| Linear Model (asin) | | ✓ | | ✓ | | ✓ | ✓ | ✓ |
| Binomial | ✓ | | | | | | | ✓ |
| Beta-binomial | | | | | ✓ | | | |
| NB | ✓ | ✓ | ✓ | ✓ | ✓ | | | ✓ |
| Poisson | ✓ | | ✓ | ✓ | | | | ✓ |
| `scCODA` | | | ✓ | | ✓ | | | |

**Figure S1: Comparison of properties of each method.** We note that incorporation of random effects, variance partitioning analysis and multivariate testing applies to all methods using a linear model, since these approaches are quite general. They could in principle be applied to generalized linear models as well, but this would require substantial methodological development. For 'Linear model', 'frac' indicates using the cell fraction, $f$, as the response, 'log' indicates $log(f)$, 'logit' indicates $logit(f) = log(f) - log(1 - f)$, and 'asin' indicates $arcsin(\sqrt{f})$.

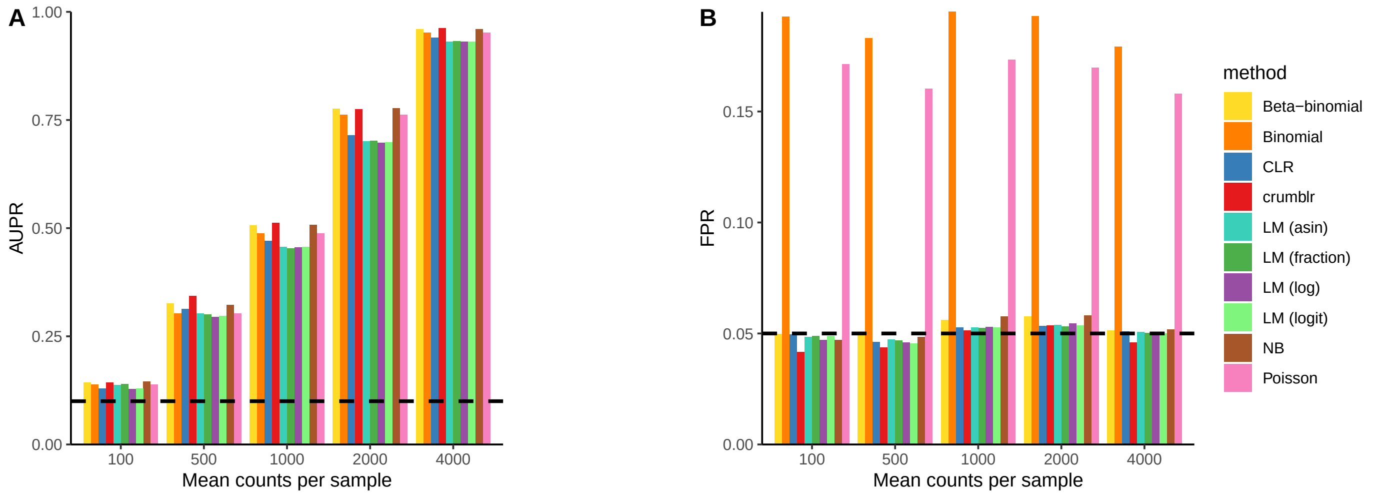


**Figure S 2: Comparing performance for increasing cell counts A)** Area under the precision recall curve. Dashed line indicates the performance of a random method. **B)** False positive rate computed as the fraction of p-values less than the target of 0.05 under a null model. Dashed line indicates the target value. Simulations were performed with 200 samples, 10 cell clusters, effect size of 0.2, mean of 2000 cells per sample, batch effect of zero, overdispersion $\tau = 10$, and 10K simulated datasets.

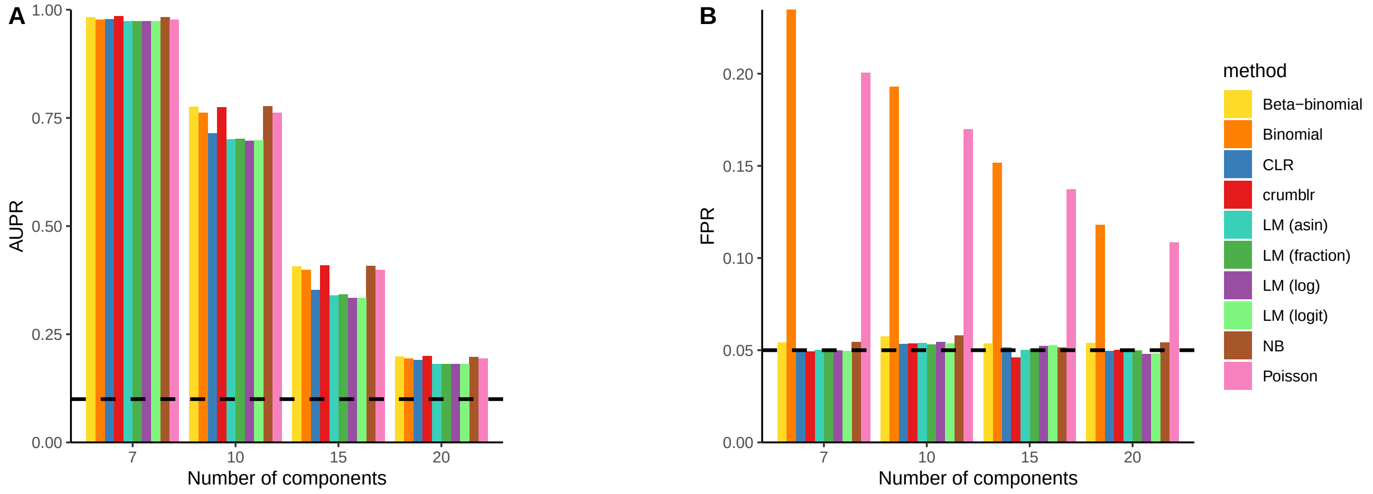


**Figure S 3: Comparing performance for increasing cell cluster components. A)** Area under the precision recall curve. Dashed line indicates the performance of a random method. **B)** False positive rate computed as the fraction of p-values less than the target of 0.05 under a null model. Dashed line indicates the target value. Simulations were performed with 200 samples, effect size of 0.2, mean of 2000 cells per sample, batch effect of zero, overdispersion $\tau = 10$, and 10K simulated datasets.

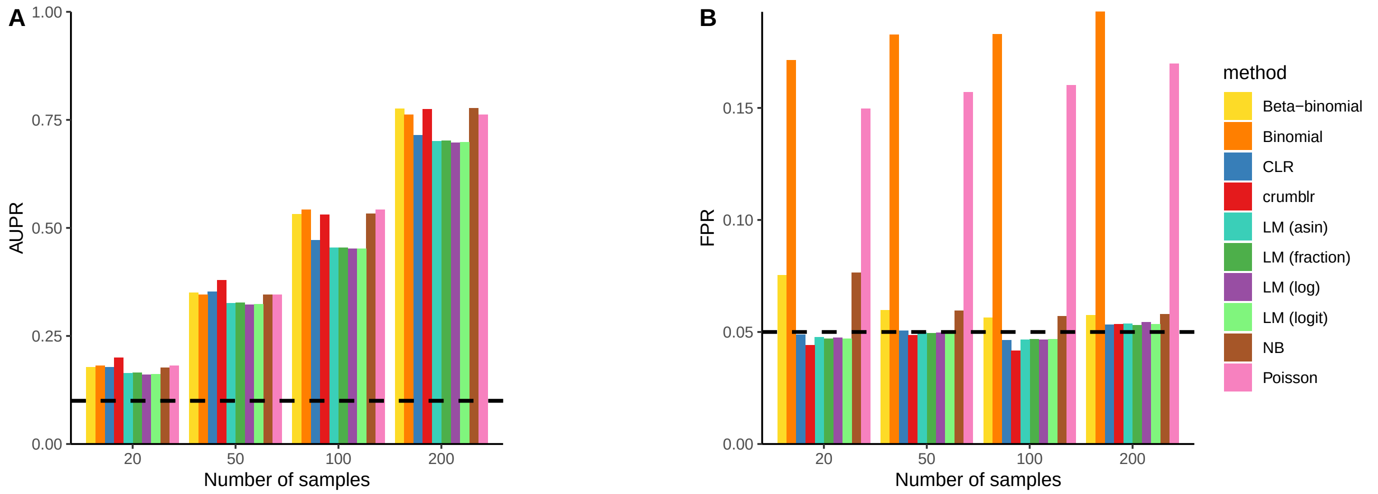


**Figure S4: Comparing performance for increasing sample size. A)** Area under the precision recall curve. Dashed line indicates the performance of a random method. **B)** False positive rate computed as the fraction of p-values less than the target of 0.05 under a null model. Dashed line indicates the target value. Simulations were performed with 10 cell clusters, effect size of 0.2, mean of 2000 cells per sample, batch effect of zero, overdispersion $\tau = 10$, and 10K simulated datasets.

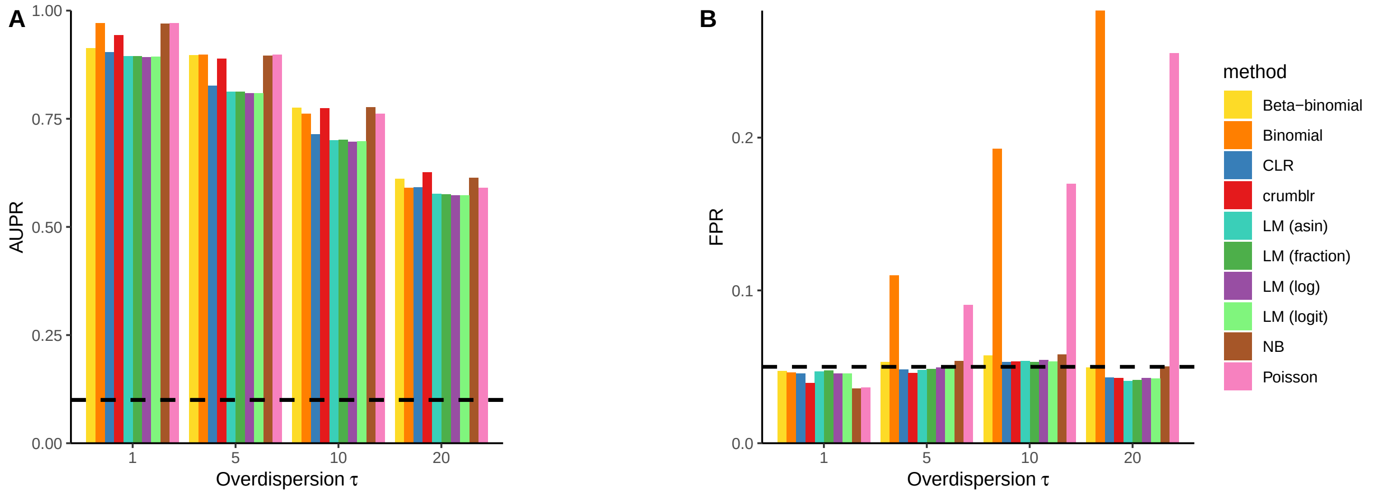


**Figure S5: Comparing performance for increasing overdispersion. A)** Area under the precision recall curve. Dashed line indicates the performance of a random method. **B)** False positive rate computed as the fraction of p-values less than the target of 0.05 under a null model. Dashed line indicates the target value. Simulations were performed with 200 samples, 10 cell clusters, effect size of 0.2, mean of 2000 cells per sample, batch effect of zero, and 10K simulated datasets.

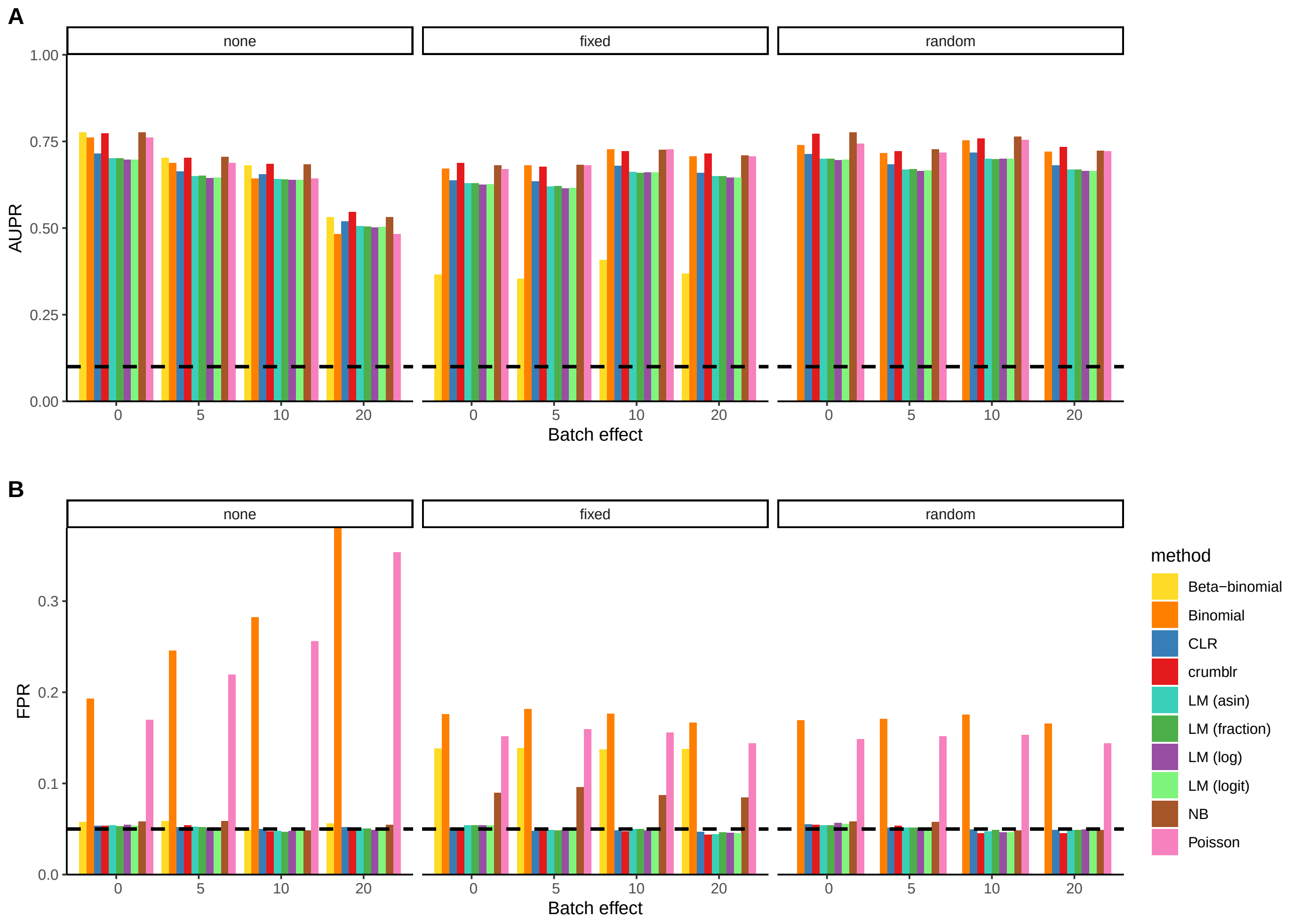


**Figure S6: Comparing performance for increasing batch effect** Results are shown when ignoring batch effect (left), and accounting for batch as a fixed effect (center) or a random effect (right). **A)** Area under the precision recall curve. Dashed line indicates the performance of a random method. **B)** False positive rate computed as the fraction of p-values less than the target of 0.05 under a null model. Dashed line indicates the target value. Simulations were performed with 200 samples, 10 cell clusters, effect size of 0.2, mean of 2000 cells per sample, overdispersion $\tau = 10$, 50 batches, and 10K simulated datasets.

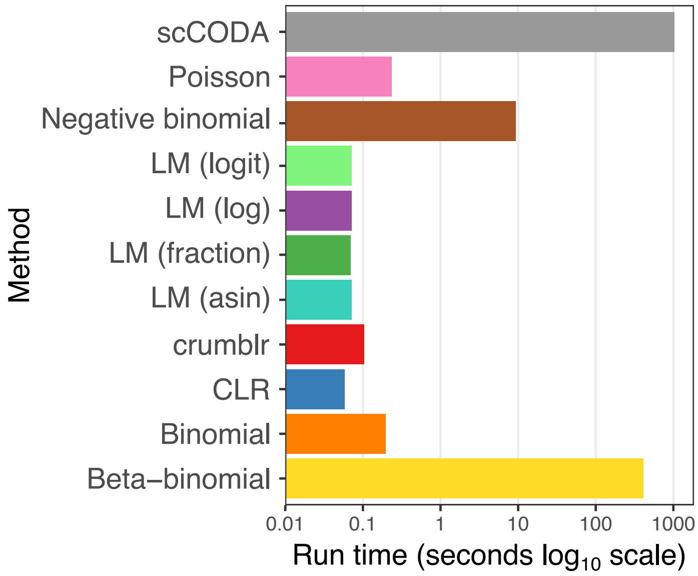


**Figure S7: Comparing compute times across methods.** Run times for fixed effect models were computed using a dataset of 500 samples, 20 clusters, 3000 cell counts per dataset, and overdispersion $\tau = 3$.

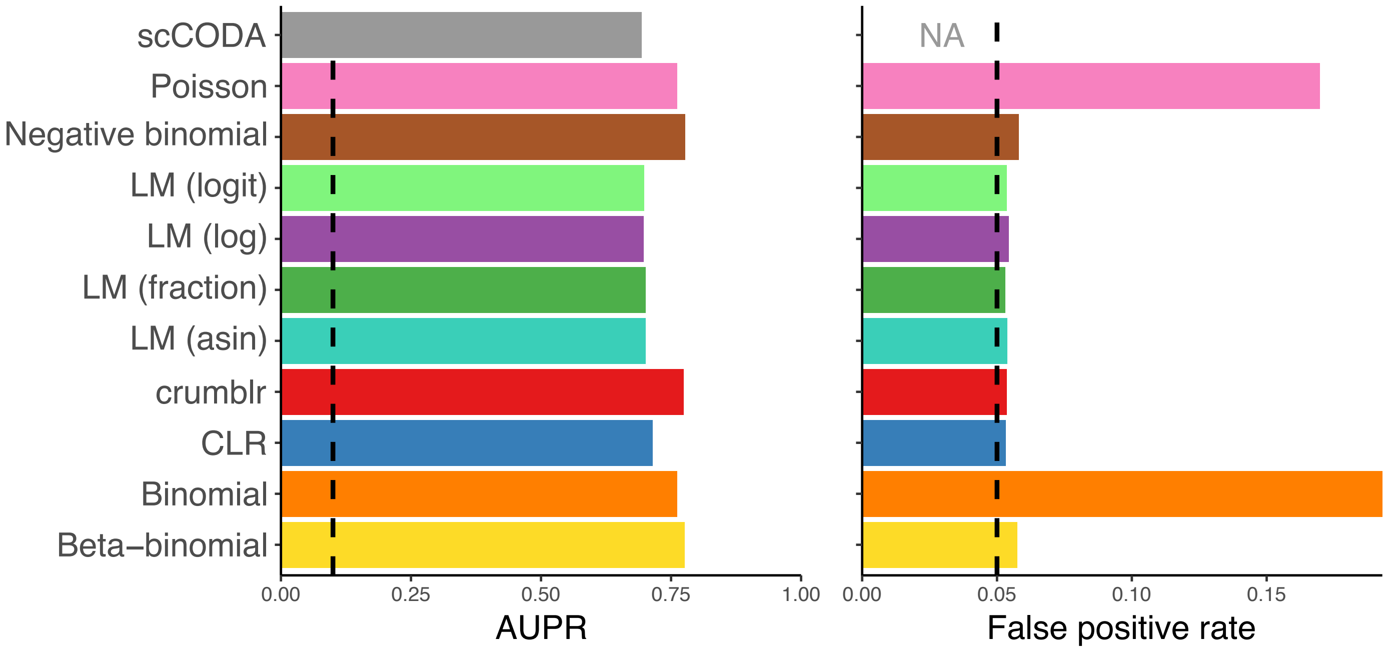


**Figure S8: Simulations using parameters identical to Figure 1A except using only 100 simulated datasets.** scCODA is not competitive with the top methods based on area under the precision recall curve (left). Because scCODA is a Bayesian model, it does not report p-values and therefore the false positive rate under the null is not applicable here. The long run time prevented a more thorough investigation of the behavior of this model.

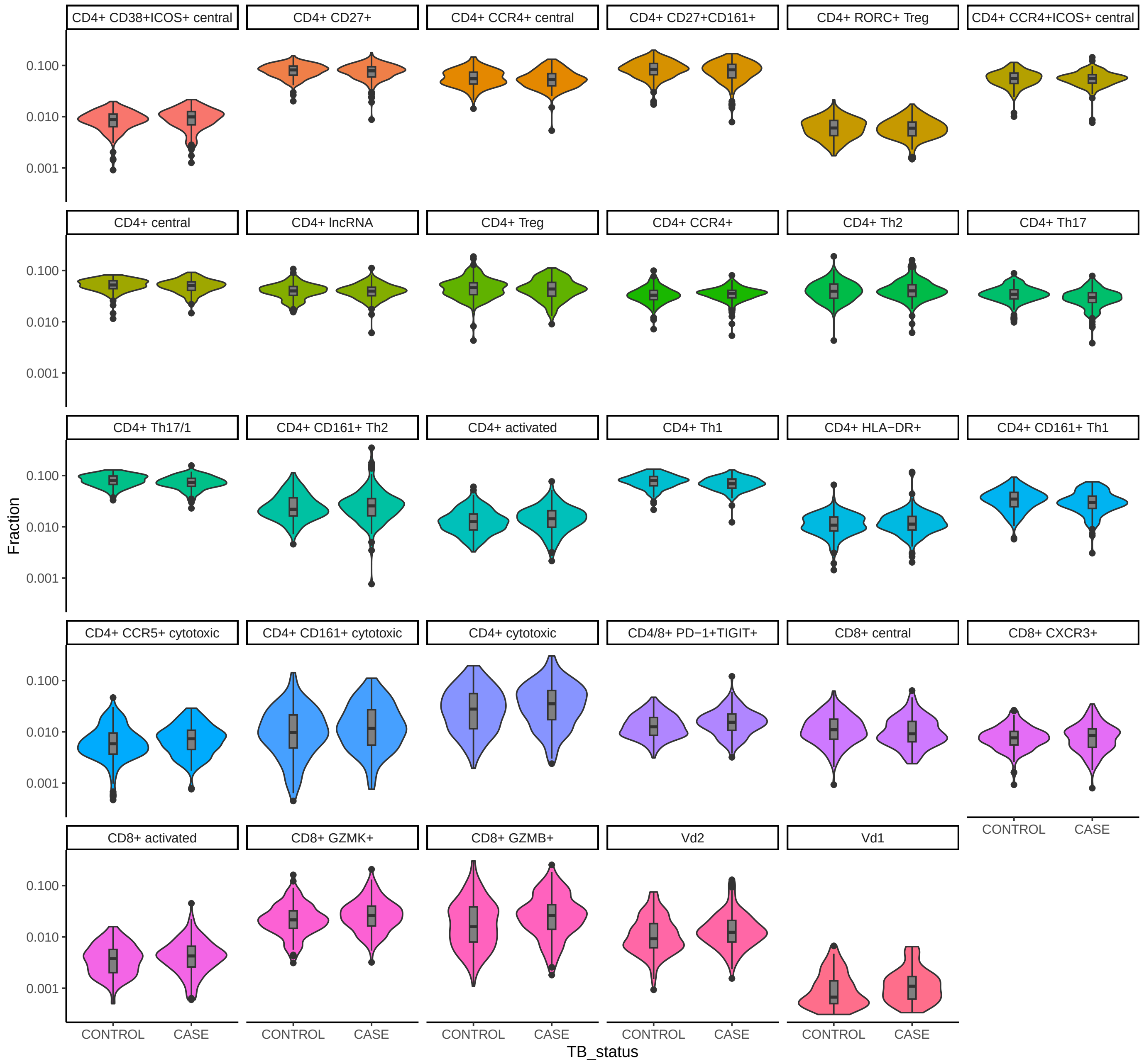


**Figure S9: Tuberculosis infection in T cell subpopulations.** Observed fractions for each cell subpopulation stratified by case vs control status.

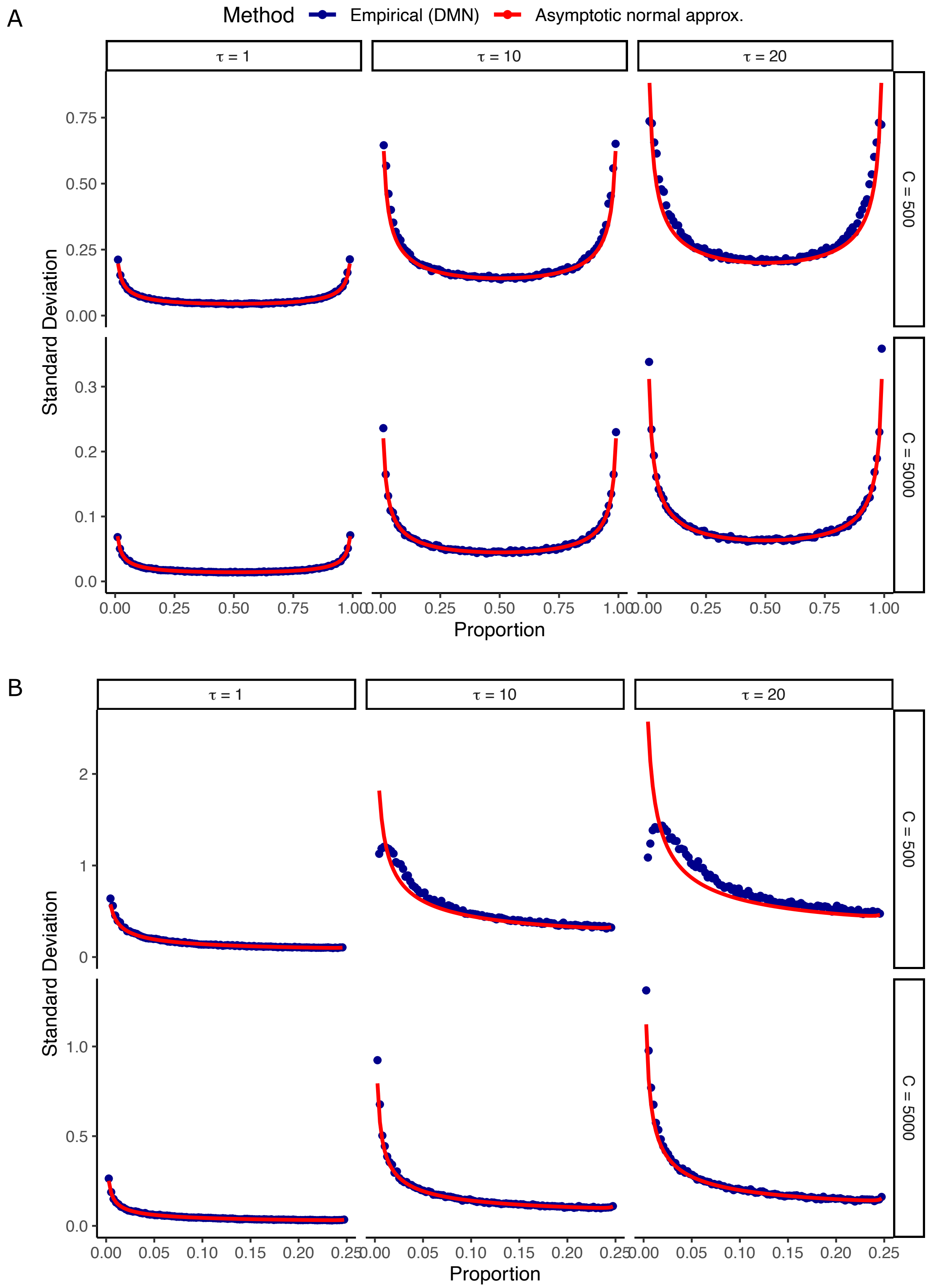


**Figure S 10: Performance of asymptotic normal approximation.** Standard deviation of the CLR transformed proportions from 1000 samples drawn from a Dirichlet multinomial model (blue points) are compared to the normal approximation developed here (red line) when at least 2 counts are observed. Data was simulated using an overdispersion $\tau \in \{1, 10, 20\}$ and total counts $C \in \{500, 5000\}$. Results are shown for **A)** $D = 2$ categories and **B)** $D = 15$ categories.

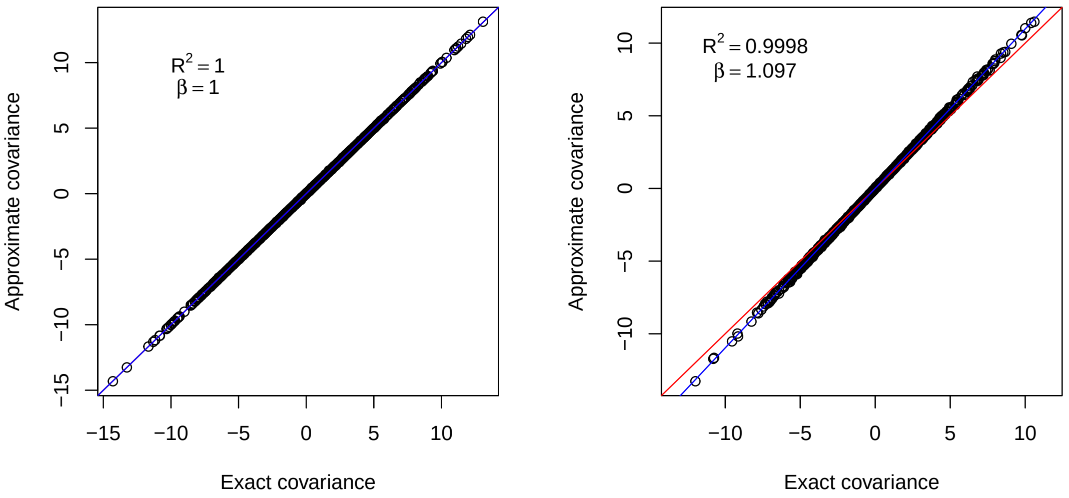


**Figure S11: Comparison of variance methods.** Compare exact versus approximate methods to compute covariance for a fixed effects model using 200 samples and 1000 simulations. Left: weights are shared across responses. Right: weights vary across responses. Red line is diagonal, blue line is the least squares fit with coefficient $\beta$.

### Supplementary Methods

#### 1.1 Count ratio uncertainty modeling with linear regression

Here we propose count ratio uncertainty modeling based linear regression (`crumblr`). When considering count ratios for multiple variables, it is desirable to compare all variables to the same baseline. Selecting the last variable as the baseline is simple, but the count ratios will depend on the ordering of the variables. Instead, Aitchison (1986) proposes the centered log ratio (clr) using the geometric mean of the counts across all variables as the common baseline:

$$\text{clr}(p_i) \quad = \quad \log(p_i) - \frac{1}{D}\sum_{j=1}^{D}\log(p_j) \tag{1}$$

There are two sources of heteroskedasticity 1) for proportions, the sample variance is related to the value. This is clearly seen with the beta distribution. 2) Due to different count totals.

For $D$ variables, these clr transform spans a $D-1$ dimensional space. This degenerate set of variables can be problematic in regression modeling. To address this Egozcue et al. (2003) proposed the isometric log ratio (ilr) to transform the clr transformed variables into $D-1$ variables. Importantly, the transformation between clr and ilr coordinates is a linear operation.

Regression models fit using clr transformed variables as responses are easily interpretable since each clr variable corresponds to an original variable. Conversely, ilr transformed variables have nice mathematical properties, but are challenging to interpret because they are linear combinations of multiple clr variables.

It is common to include clr and ilr transformed variables as responses in regression models. However, standard application does not account for the substantial difference in measurement precision across observations. The standard assumption of equal measurement precision in regression models can cause loss of power and poor control of the false positive rate.

Here, we take advantage of the interpretability of the clr transform and the mathematical properties of the ilr transform to explicitly model measurement uncertainty in observed counts with precision weights.

##### 1.1.1 Asymptotic normal approximation of multinomial counts

Let $\mathbf{p}$ be the true vector of proportions for $D$ components with $p_i$ being the true proportion for the $i^{th}$ component. Let $\mathbf{c}$ be the vector of observed counts drawn based on a multinomial distribution with $n$ total counts. The expected number of counts is clearly $E[\mathbf{c}] = n\mathbf{p}$, and the maximum likelihood estimate of the proportions is $\hat{\mathbf{p}} = \mathbf{c}/n$, and the variance of the counts is $n\left[\text{diag}(\mathbf{p}) - \mathbf{p}\mathbf{p}^T\right]$. Based asymptotic properties of multinomial distribution, for large $n$ the distribution of the estimated proportions converges to a multivariate normal distribution

$$\hat{\mathbf{p}} \xrightarrow{d} \mathcal{N}\left(\mathbf{p}, \frac{1}{n}\left[\text{diag}(\mathbf{p}) - \mathbf{p}\mathbf{p}^T\right]\right). \tag{2}$$

Egozcue et al. (2020) and Graffelman et al. (2015) propose an asymptotic normal approximation of the ilr transformed components. The ilr transform is defined by a linear transformation

$$\text{ilr}(\mathbf{p}) = V^T \text{clr}(\mathbf{p}) \tag{3}$$

where $V$ is a $D \times D - 1$ matrix with orthogonal columns that projects the clr-transformed components into a lower dimensional space (Egozcue et al., 2003). By construction $V$ satisfies properties

$$\begin{aligned} V^T V &= I_{D-1} & (4)\\ V V^T &= I_D - \frac{1}{D}\mathbf{1}\mathbf{1}^T & (5)\\ V^+ &= V^T & (6) \end{aligned}$$

where $V^+$ indicates the Moore–Penrose inverse. It follows that the inverse projection from ilr coordinates back to clr coordinates is

$$\text{clr}(\mathbf{p}) = V\,\text{ilr}(\mathbf{p}) \tag{7}$$

Egozcue et al. (2020) and Graffelman et al. (2015) use the delta method to obtain the asymptotic normal distribution of the ilr transformed estimated proportions:

$$\text{ilr}(\mathbf{p}) \to \mathcal{N}\left(\text{ilr}(n\mathbf{p}), \frac{1}{n}V^T \text{diag}(1/\mathbf{p})V\right). \tag{8}$$

Given this variance matrix in ilr space, we derive the variance-covariance matrix in clr space using the projection matrices mapping between the two spaces:

$$\begin{aligned} \text{var}[\text{clr}(\mathbf{p})] &= V\,\text{var}[\text{ilr}(\mathbf{p})]V^T & (9)\\ &= \frac{1}{n} V V^T \text{diag}(1/\mathbf{p}) V V^T & (10)\\ &= \frac{1}{n}\left(I_D - \frac{1}{D}\mathbf{1}\mathbf{1}^T\right)\text{diag}(1/\mathbf{p})\left(I_D - \frac{1}{D}\mathbf{1}\mathbf{1}^T\right)^T & (11)\\ &= \frac{1}{n}\left(\text{diag}(1/\sqrt{\mathbf{p}}) - \frac{1}{D}\mathbf{1}\mathbf{1}^T\text{diag}(1/\sqrt{\mathbf{p}})\right)\left(\text{diag}(1/\sqrt{\mathbf{p}}) - \frac{1}{D}\mathbf{1}\mathbf{1}^T\text{diag}(1/\sqrt{\mathbf{p}})\right)^T & (12)\\ &= \frac{1}{n}\left[\text{diag}(1/\mathbf{p}) - \frac{2}{D}\text{diag}(1/\sqrt{\mathbf{p}})\mathbf{1}\mathbf{1}^T\text{diag}(1/\sqrt{\mathbf{p}}) + \frac{1}{D^2}\mathbf{1}\mathbf{1}^T\text{diag}(1/\mathbf{p})\mathbf{1}\mathbf{1}^T\right] & (13) \end{aligned}$$

This describes the full $D \times D$ asymptotic variance-covariance matrix of the clr-transformed counts. The sampling variance of the $i^{th}$ component is

$$\text{var}[\text{clr}_i(\mathbf{p})] = \frac{1}{n}\left[\frac{1}{p_i} - \frac{2}{Dp_i} + \frac{1}{D^2}\sum_{j=1}^{D}\frac{1}{p_j}\right] \tag{14}$$

For observed data, the maximum likelihood estimate of the proportion can be substituted for the true value to obtain an estimate of the variance

$$\widehat{\text{var}[\text{clr}_i(\mathbf{p})]} = \frac{1}{n}\left[\frac{1}{\hat{p}_i} - \frac{2}{D\hat{p}_i} + \frac{1}{D^2}\sum_{j=1}^{D}\frac{1}{\hat{p}_j}\right] \tag{15}$$

Note that the variance is a function of the total number of counts, number of components, and the proportions for each component. This value is easy to compute for each component.

##### 1.1.2 Asymptotic normal approximation to Dirichlet Multinomial

The Dirichlet Multinomial (DMN) distribution is a more flexible model of multi-class count data that generalizes the multinomial distribution. While the multinomial distribution assumes the underlying proportions are fixed, in the DMN the proportions are random variables drawn from a Dirichlet distribution. Counts from a DMN model can be drawn according to

$$\mathbf{x} \sim \textit{Multinomial}(n, \mathbf{p}) \tag{16}$$

$$\mathbf{p} \sim \textit{Dirichlet}(\boldsymbol{\alpha}) \tag{17}$$

where $\boldsymbol{\alpha}$ is the vector of parameters for Dirichlet distribution so that the expected proportion of category $i$ is $\alpha_i/\alpha_0$ where $\alpha_0 = \sum_k \alpha_k$. The sampling variance of observed counts is now

$$\text{var}(\mathbf{c}) = \tau\mathbf{n}\left[\text{diag}(\mathbf{p}) - \mathbf{p}\mathbf{p}^{\mathbf{T}}\right]. \tag{18}$$

where $\tau = \frac{n+\alpha_0}{1+\alpha_0}$ is the overdispersion parameter. Setting $\tau = 1$ produces the multinomial distribution as a special case.

Since the overdispersion $\tau$ is a scalar parameter shared by all categories, the derivation from the previous section can be adapted to show that the proportions are asymptotically normal according to

$$\hat{\mathbf{p}} \xrightarrow{d} \mathcal{N}\left(\mathbf{p}, \frac{\tau}{n}\left[\text{diag}(\mathbf{p}) - \mathbf{p}\mathbf{p}^{T}\right]\right), \tag{19}$$

so that the asymptotic variance-covariance matrix of the clr-transformed proportions is

$$\widehat{\text{var}[\text{clr}_i(\mathbf{p})]} = \frac{\tau}{n}\left[\frac{1}{\hat{p}_i} - \frac{2}{D\hat{p}_i} + \frac{1}{D^2}\sum_{j=1}^{D}\frac{1}{\hat{p}_j}\right]. \tag{20}$$

This approximation is quite accurate even for small fractions when a sufficient number of total cells are observed (**Figure S10**).

#### 1.2 Multivariate testing combining multiple cell type components

Consider a multivariate regression with $n$ samples, $c$ variables, and $m$ response values using design matrix $X_{n\times c}$ and responses stored as column in the matrix $Y_{n\times m}$. When the regression weights are all equal to one, standard results give estimated coefficients

$$\hat{B} = \left(X^T X\right)^{-1} X^T Y. \tag{21}$$

Letting the error covariance be $C = R^T R/d$ with $d$ the residual degrees of freedom, the covariance matrix of the coefficient estimates is

$$\text{var}(\text{vec}(\hat{B})) = C \otimes \left(X^T X\right)^{-1} \tag{22}$$

where the vec(.) operator converts a matrix to a column vector, and $\otimes$ is the Kronecker product. Examining covariance between coefficients from response $i$ and $j$ gives

$$\begin{aligned} \text{cov}\left(\hat{\beta}_i, \hat{\beta}_j\right) &= C_{i,j}\left(X^T X\right)^{-1} & (23)\\ &= \text{cor}(r_i, r_j)\left(X^T X\right)^{-1}\sigma_i\sigma_j. & (24)\end{aligned}$$

where $C = r_i^T r_j/d$, and $\hat{\beta}_i$ is the $i^{th}$ column of $\hat{B}$.

##### 1.2.1 Weighted linear model

When weights are not equal to 1 and vary for each response, the coefficient estimates and their covariance have a more complicated form. Letting $w^{(i)}$ be the vector of weights for response $i$, $W^{(i)} = \text{diag}\left(w^{(i)}\right)$, and $y_i$ be the vector storing response $i$,

$$\hat{\beta}_i = \left(X^T W^{(i)} X\right)^{-1} X^T W^{(i)} y_i \tag{25}$$

Creating a weighted version of any matrix or vector according to $\tilde{M}_i = M\text{diag}\left(\sqrt{w^{(i)}}\right)$, simplifies the notation so that the estimated coefficients are

$$\hat{\beta}_i = \left(\tilde{X}_i^T \tilde{X}_i\right)^{-1} \tilde{X}_i^T \tilde{y}_i, \tag{26}$$

and the covariance between coefficients estimates is now

$$\text{cov}\left(\hat{\beta}_i, \hat{\beta}_j\right) = \tilde{C}_{i,j}\left(\tilde{X}_i^T \tilde{X}_i\right)^{-1}\left(\tilde{X}_i^T \tilde{X}_j\right)\left(\tilde{X}_j^T \tilde{X}_j\right)^{-1}. \tag{27}$$

where $\tilde{C} = \tilde{r}_i^T \tilde{r}_j/d$. When all weights equal 1, these reduce to the standard equations (21) and (21).

##### 1.2.2 Weighted linear mixed model

For the linear mixed model, closed form estimates don't exist. Each coefficient vector is estimated using numerical optimization in `lme4` (Bates et al., 2015) with the covariance matrix between coefficients for response $i$ indicated by $\hat{\Sigma}_i$. Using the form of equation (24), the covariance in coefficient estimates between responses $i$ and $j$ can be stated in terms of the covariance for each response and the correlation between residuals when all weights are 1. The covariance between coefficients estimates for fixed effects in the linear mixed model is approximated as

$$\text{cov}\left(\hat{\beta}_i, \hat{\beta}_j\right) = \text{cor}(\tilde{r}_i, \tilde{r}_j)\hat{\Sigma}_i \left(\hat{\Sigma}_i^{-\frac{1}{2}} \hat{\Sigma}_j^{-\frac{1}{2}}\right) \hat{\Sigma}_j \tag{28}$$

When $i = j$, this formula reduces to the $\hat{\Sigma}_i$. In the case of linear regression, when the weights vary across samples but are shared across responses, this method is equivalent to equation (24). In the case of general weights, simulations indicate that approximation in equation in (28) produces accurate estimates of the covariance in coefficient estimates across responses (**Figure S11**).

##### 1.2.3 Fixed effect hypothesis testing accounting for covariance

Letting $B_{k,.}$ indicate the vector of estimates for coefficient $k$ across responses, $\Omega$ be the covariance matrix of these coefficient estimates with values computed by equations (27) or (28). A multivariate hypothesis test is performed for coefficient $k$ across multiple responses using a fixed effects meta analysis designed for correlated coefficient estimates (Han et al., 2016, Lee et al., 2017, Lin and Sullivan, 2009). Under this model the test statistic

$$\frac{\left(\mathbf{1}^T \Omega^{-1} B_{k,.}\right)^2}{\mathbf{1}^T \Omega^{-1} \mathbf{1}}, \tag{29}$$

where $\mathbf{1}$ is a vector of 1's, is distributed as a $\chi^2_1$ under the null. When $\Omega$ is diagonal, so there is no covariance between coefficient estimates, this reduce to standard fixed effects meta-analysis (Han et al., 2016).

#### 1.3 Simulations

##### 1.3.1 Univariate hypothesis testing

Counts were simulated from a Dirichlet-multinomial with a specified sample size, number of classes, overdispersion, effect size of the trait variable, and size of the batch effect. The trait of interest was normally distributed and samples were randomly assigned discrete batches. The coefficients for the effect of each batch were normally distributed with a mean of zero and a variance of $\gamma \in (0, 0.5, 1, 3)$.

The linear predictor was created by multiplying the design matrix storing the trait and batch variables by the coefficient vector. The linear predictor for each sample was then exponentiated to obtain the fractions for each cell component. For each sample, the total number cell counts was drawn from a negative binomial with specified mean and $\theta = 3$ to give substantial variation in total cell counts as seen in real data. Counts were drawn from a Dirichlet-multinomial based on the probabilities, total count number per sample, and overdispersion.

##### 1.3.2 Multivariate hypothesis testing

Results are shown for the fixed effect model incorporating correlation between test statistics (Lin and Sullivan, 2009), Fisher's method for combining p-values across tests ($-2\sum_{i=1}^{k} \log p_i \sim \chi^2_{2k}$ for $k$ p-values), and Sidak's method for computing the minimum p-value corrected for multiple testing ($p_{\text{Sidak}} = 1 - (1 - \min(p))^k$). Simulations were performed 100k times with with 200 samples, 2 to 12 responses, heteroskedastic measurement error, and an error correlation between response variables of 0.8. 20k simulations used an effect sizes of 0.8 shared across responses, while the rest were simulated under the null model of no effect.

#### 2 Implementation of other methods for comparison

Hypothesis testing for differential cell composition was performed with a range of existing methods that either model the cell fraction or directly modeled the number of observed counts. Let `y` be a vector storing the observed counts for a given cell type across all samples, `C` be a vector storing the total number of cells observed for each sample, and `x` be the variable of interest for hypothesis testing. Most models were fit in `R` as described in **Table S1** for fixed effects and **Table S2** for mixed effect modeling of batches. We note that the beta-binomial model does not supported mixed effects.

The centered log ratio (CLR) method was fit by performing the CLR transform using `clr()` in the `compositions` package available on CRAN and then fitting the regression models for each cell type using `dream()` in the `variancePartition` package (Hoffman and Roussos, 2021, Hoffman and Schadt, 2016). Transformations involving log, logit and asin were computed this way too. The `crumblr()` function from the `crumblr` package was used to transform the count data and estimate the measurement precision of the transformed abundance values before being fed into `dream()`.

`scCODA` is a Bayesian method implemented in python (Büttner et al., 2021) and was called with default parameters using the observed cell counts and design matrix.

### Supplementary Tables

| Method | Function | Formula | Arguments |
|---:|---|---|---|
| Poisson | `glm()` | `y ~ offset(log(C)) + x` | `family=poisson()` |
| Negative binomial | `glm.nb()` | `y ~ offset(log(C)) + x` | |
| LM (fraction) | `lm()` | `y / C ~ x` | |
| LM (log) | `lm()` | `log(y / C) ~ x` | |
| LM (logit) | `lm()` | `logit(y / C) ~ x` | |
| LM (asin) | `lm()` | `asin(sqrt(y / C)) ~ x` | |
| Binomial | `glm()` | `cbind(y, C - y) ~ x` | `family=binomial()` |
| Beta-binomial | `betabin()` | `cbind(y, C - y) ~ x` | |

**Table S1: Implementation of each method for fixed effects models**

<table>
<tr><th>Method</th><th>Function</th><th>Formula</th><th>Arguments</th></tr>
<tr><td>Poisson</td><td>glmer()</td><td>y ~ offset(log(C)) + (1|Batch) + x</td><td>family=poisson()</td></tr>
<tr><td>Negative binomial</td><td>glmer.nb()</td><td>y ~ offset(log(C)) + (1|Batch)+ x</td><td></td></tr>
<tr><td>LM (fraction)</td><td>lmer()</td><td>y / C ~ (1|Batch) + x</td><td></td></tr>
<tr><td>LM (log)</td><td>lmer()</td><td>log(y / C) ~ (1|Batch) + x</td><td></td></tr>
<tr><td>LM (logit)</td><td>lmer()</td><td>logit(y / C) ~ (1|Batch) + x</td><td></td></tr>
<tr><td>LM (asin)</td><td>lmer()</td><td>asin(sqrt(y / C)) ~ (1|Batch) + x</td><td></td></tr>
<tr><td>Binomial</td><td>glmer()</td><td>cbind(y, C - y) ~ (1|Batch) + x</td><td>family=binomial()</td></tr>
</table>

**Table S2: Implementation of each method for mixed effects models**
